# PanEffect: A pan-genome visualization tool for variant effects in maize

**DOI:** 10.1101/2023.09.25.559155

**Authors:** Carson M Andorf, Olivia C Haley, Rita K Hayford, John L Portwood, Shatabdi Sen, Ethalinda K Cannon, Jack M Gardiner, Margaret R Woodhouse

## Abstract

Understanding the effects of genetic variants is crucial for accurately predicting traits and phenotypic outcomes. Recent advances have utilized protein language models to score all possible missense variant effects at the proteome level for a single genome, but a reliable tool is needed to explore these effects at the pan-genome level. To address this gap, we introduce a new tool called PanEffect. We implemented PanEffect at MaizeGDB to enable a comprehensive examination of the potential effects of coding variants across 51 maize genomes. The tool allows users to visualize over 550 million possible amino acid substitutions in the B73 maize reference genome and also to observe the effects of the 2.3 million natural variations in the maize pan-genome. Each variant effect score, calculated from the Evolutionary Scale Modeling (ESM) protein language model, shows the log-likelihood ratio difference between B73 and all variants in the pan-genome. These scores are shown using heatmaps spanning benign outcomes to strong phenotypic consequences. Additionally, PanEffect displays secondary structures and functional domains along with the variant effects, offering additional functional and structural context. Using PanEffect, researchers now have a platform to explore protein variants and identify genetic targets for crop enhancement.

Availability and implementation: The PanEffect code is freely available on GitHub (https://github.com/Maize-Genetics-and-Genomics-Database/PanEffect). A maize implementation of PanEffect and underlying datasets are available at MaizeGDB (https://www.maizegdb.org/effect/maize/).

## 1 Introduction

Genetic variations play an important role in determining phenotypic outcomes in plants, influencing traits such as yield, disease resistance, and environmental adaptability. The challenges of food security, changing climates, and evolving pathogens have made understanding the functional consequences of genetic variation increasingly critical. In crops, genetic variation studies have contributed to the development of drought-resistant and disease-resistant cultivars (Cooper and Messina, 2023). These investigations, however, often demand considerable time and effort. Thus, a method that efficiently links expansive genetic datasets and functional annotations to phenotypic differences is needed to facilitate these efforts. We introduce PanEffect, an AI-driven platform tailored for the maize genome that offers insights into the effects of genetic variants in maize. This tool allows researchers to access detailed maize gene summaries and visualize variant effects in the B73 line and across the maize pan-genome.

Previous efforts designed to predict genome-wide variant effects (Bernhofer *et al*., 2021; Sun and Shen, 2023; W. Lin *et al*., 2023; Ramakrishnan *et al*., 2023; Wagih *et al*., 2018; McLaren *et al*., 2016; Mahmood *et al*., 2017; Fowler *et al*., 2023; Gray *et al*., 2018; Abakarova *et al*., 2023; Cheng *et al*., 2023; Brandes *et al*., 2023) set the stage for the development of platforms like PanEffect. For example, the AlphaMissense resource (Cheng *et al*., 2023) combined structural and evolutionary information to rate all possible missense mutations in the human genome as either benign or likely pathogenic. In a second example, the GEMME framework (Laine *et al*., 2019) was used as an alignment-based strategy for predicting mutational outcomes across numerous protein families by assessing 1.5 million missense variants, highlighting the feasibility of high-quality and computationally efficient mutational landscape predictions for complete proteomes (including the human proteome) (Abakarova *et al*., 2023). In a third example, a workflow called esm-variants used a large-scale protein language model to predict the effects of all potential missense variants in the human genome, achieving high performance in variant classification and demonstrating the applicability of protein language models in predicting variant effects for various coding variants (Brandes *et al*., 2023).

While these projects focus on identifying all possible coding substitutions for a reference genome, PanEffect is unique by combining reference-genome coding substitutions with additional tools and datasets to show the predicted variant effects across the natural variation in a pan-genome, as shown in our MaizeGDB instance with 51 maize genomes. It integrates over 550 million possible missense mutations in the maize reference genome with protein annotations from sources such as UniProt (UniProt: the universal protein knowledgebase in 2021, 2021), Pfam (Mistry *et al*., 2021), and trait data from genome-wide association studies hosted at MaizeGDB. With the capability to predict the phenotypic outcome of specific genetic changes, PanEffect will help bridge the gap between genetic information and observable traits to help researchers develop new maize varieties or obtain a clearer understanding of genetic variations across the maize pan-genome.

## 2 Application description

The PanEffect instance can be implemented for any proteome, but the instance described here is customized for proteins in the B73 RefGen_v5 (Zm-B73-REFERENCE-NAM-5.0) maize reference genome and is available in the Maize Genetics and Genomics Database (MaizeGDB) (Woodhouse *et al*., 2021; Portwood *et al*., 2019; Cannon *et al*., 2011). The web interface has five components: Gene Summary, Variant Effects in B73, Variant Effects across the pan-genome, Search, and Help. Each component executes fully client-side using JavaScript to manage the front-end visualization. There are no backend databases or server-side programming languages required. The overviews of the B73 and pan-genome views are shown in **Figures 1** and **2**, respectively. PanEffect is organized in a single HTML index file, five JavaScript files, and a CSS file. The index file contains the framework for the displays. The JavaScript files manage the data loading (main.js), reference genome visualization (genome.js), pan-genome visualization (pan.js), webpage layout (dom.js), and support functions (support.js). The CSS file contains the styles and formatting of the tool. A description of each of the five major components, provided as tabs in PanEffect, is provided below.

**Figure 1:**
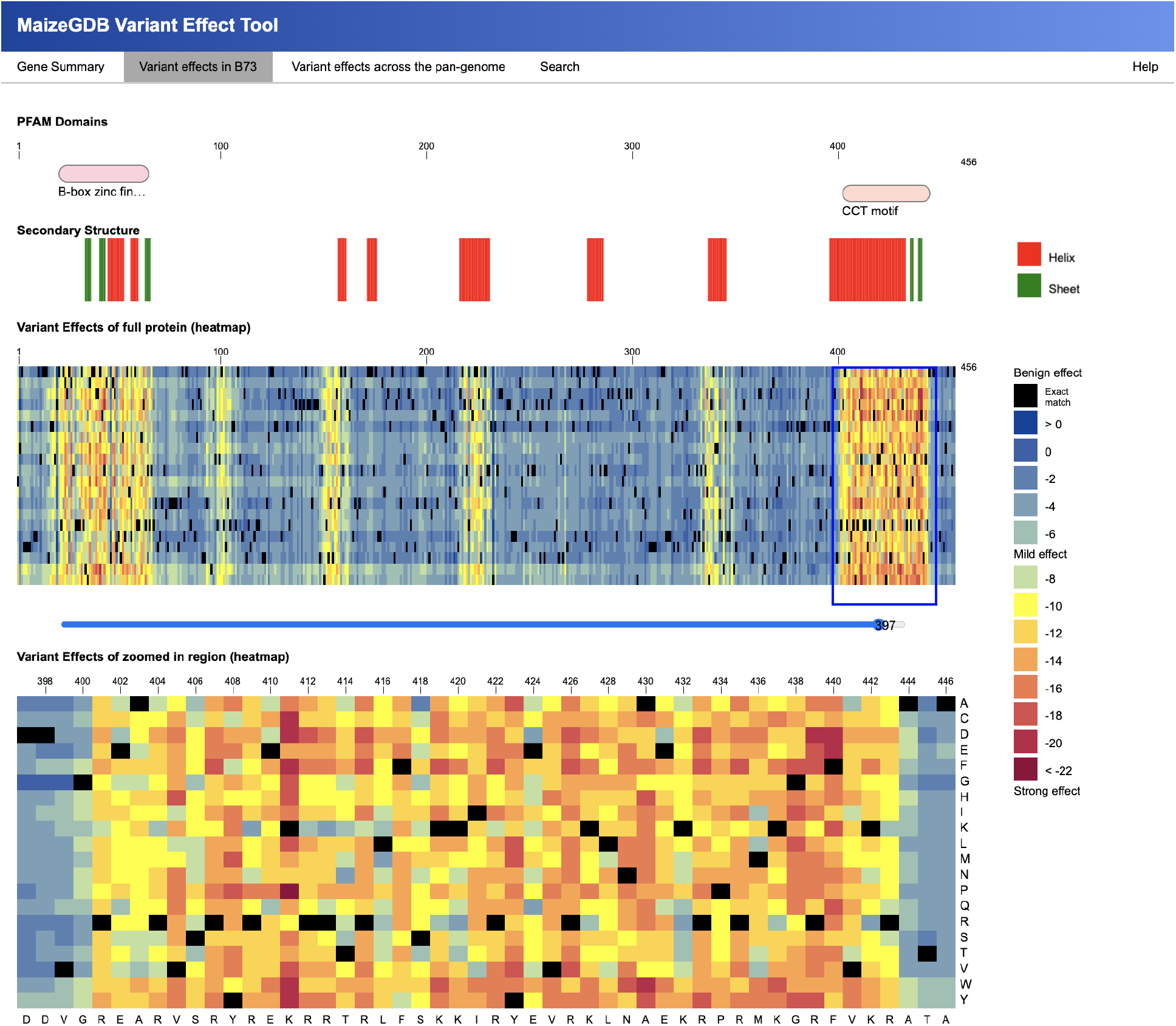
Overview of the ‘Variant effects in B73’ view. A snapshot of the PanEffect website for the ‘Variant effects in B73’ view for gene model Zm00001eb379950 (gene symbol: col4; gene name: C2C2-CO-like-transcription factor 4). Top to bottom: Pfam domains; predicted secondary structures; heatmap of the variant effects scores for all possible amino acid substitutions; navigation scrollbar; and a zoomed-in view of the heatmap around the CCT motif. The scrollbar in the center of the page allows a user to control which portion of the protein to view in the zoomed-in display. The variant effects are color-coded based on the level of potential phenotypic effect. This snapshot shows a common example of potential strong phenotypic effects in regions corresponding to functional domains and predicted secondary structures.

**Figure 2:**
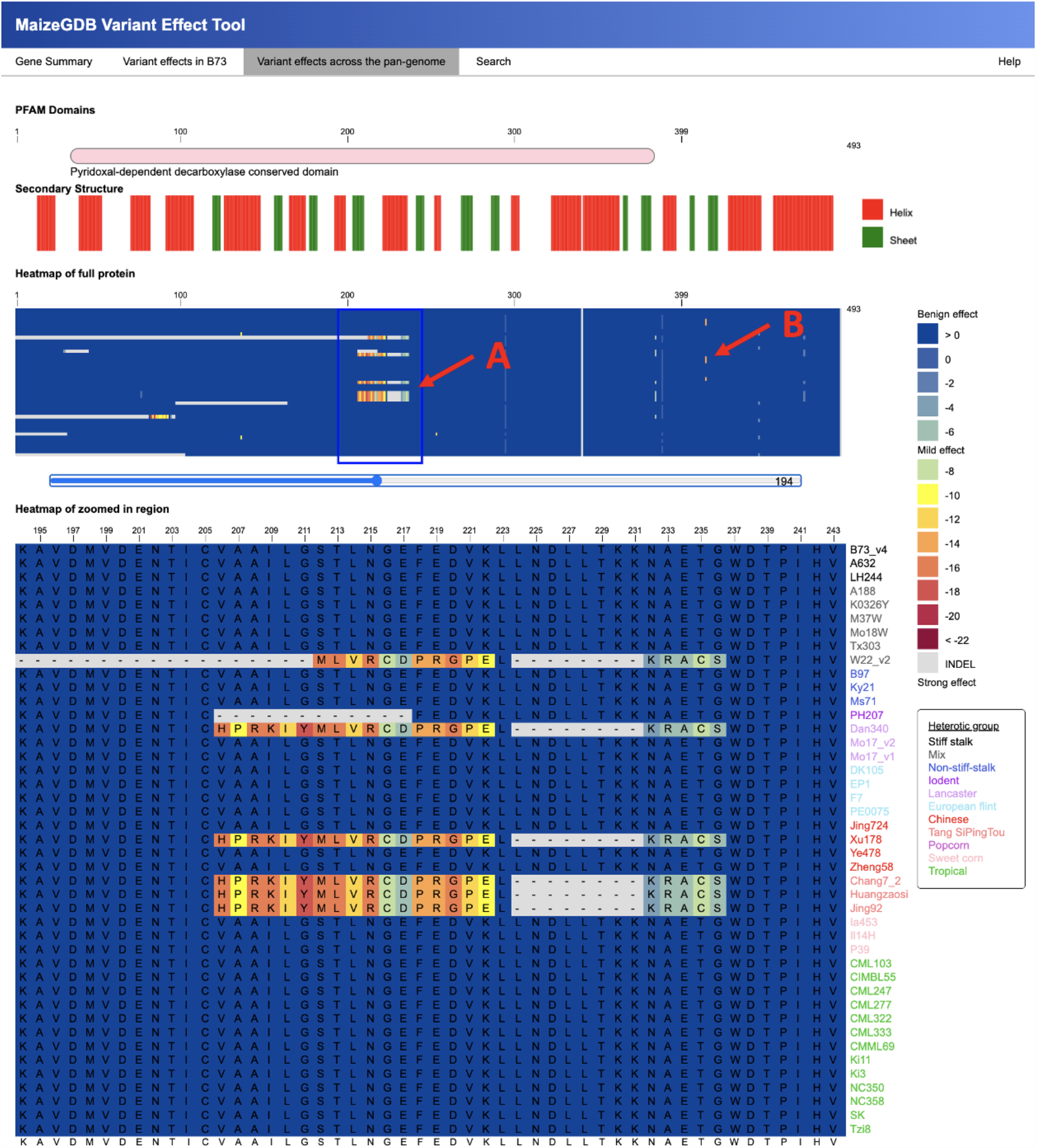
Overview of the ‘Variant effects across the pan-genome’ view. The figure shows a snapshot of the ‘Variant effects across the pan-genome’ view for gene model Zm00001eb055490 (Glutamate decarboxylase). Top to bottom: Pfam domains; predicted secondary structures; heatmap of variant effects scores of naturally occurring variations in the maize pan-genome; and a zoomed-in region showing a region where a few proteins have high variant effect scores. Each amino acid variant is color-coded based on the level of potential phenotypic effect, and each genome is color-coded based on the heterotic group. Insertions and deletions between the reference and target proteins are shown as gray vertical and horizontal regions on the display and labeled with a ‘-’. This example shows two areas of strong effect: Arrow A highlights a stretch of strong variation starting at position 206 overlapping the Pfam domain found in five maize lines used in Chinese breeding programs and the W22 genome, but are not present in the other heterotic groups. Arrow B points to a region where a histidine converted to a proline is predicted to have a strong effect at position 413 for the genomes A188, K0326Y, Mo17, and Jing724.

### Gene Summary

The gene summary section provides a detailed look at annotations and valuable information regarding gene models and proteins. The gene summary is customizable for each instance of PanEffect. The MaizeGDB instance includes gene and gene model annotations from MaizeGDB, protein annotations from UniProt (UniProt: the universal protein knowledgebase in 2021, 2021), links to 3D structure tools (Woodhouse *et al*., 2023), and trait data from three collections of genome-wide association studies (Wallace *et al*., 2014; Tian *et al*., 2020; Li *et al*., 2022).

### Variant Effects in B73

The reference genome variant effects view provides a visualization of the variant effects of all possible amino acid substitutions in the B73 maize line. This section has two heatmaps, one providing a broad overview and another providing a detailed view. The columns of the heatmap represent each position of the reference protein, and the rows represent the 20 possible amino acid substitutions. The colors of the cells within the heatmap shift from blue (benign outcomes) to red (strong phenotypic impact). Hovering over a cell provides additional information about the possible substitution including position, substitution, and score. A slider bar in the center of the page controls which portion of the protein is shown on the zoomed-in heatmap. Two tracks above the heat show the locations of Pfam domains and predicted secondary structures which further provide context regarding the functional and structural roles of each amino acid and its possible role in the consequence of amino acid substitutions.

### Variant Effects across the Pan-genome

The pan-genome variant effects view also displays heatmaps with a broad and detailed view. However, this instance shows the effects of naturally occurring maize protein variations across the pan-genome. The columns of the heatmap represent each position of the protein of the reference protein. The heatmap has a row for each protein in the pan-genome that is aligned to the reference protein. Insertions and deletions in the alignments of the proteins are characterized by a ‘-’. The cells in the heatmaps range from blue (benign outcomes) to red (strong phenotypic impact) for variants within the pan-genome as compared to the reference protein. Hovering over a cell provides additional information about the known substitution including B73 position, target position, target genome, target gene model, B73 and variant amino acids, and score. This view is only available for the canonical transcript of each gene model. For B73, canonical transcripts are selected based on domain coverage, protein length, and similarity to assembled transcripts, representing a standardized or reference version of a gene’s structure (Hufford *et al*., 2021).

### Search

The search section has a search bar and a summary of the tool. The search accepts gene symbols (‘wx1’), gene models (‘Zm00001eb378140’), transcripts (‘Zm00001eb378140_T001’), and protein identifiers (‘Zm00001eb378140_P001’).

### Help

The help section offers summaries of all visualization components and includes detailed descriptions and links to the data sources, tools, downloads, and references that were utilized in the development of PanEffect. Additionally, while PanEffect uses B73 as a reference genome, it also provides links to download variant effect scores for all 51 maize genomes used in the pan-genome, totaling over 20 billion variant effect scores for maize.

### Data preparation

PanEffect integrates seven different datasets to explore the potential phenotypic consequences of missense mutations. Various tools and datasets were used to generate input files for PanEffect. This section provides a description of each of the data types and how they were generated. Full descriptions and examples are provided on GitHub. Downloads of the full datasets are available at MaizeGDB.

### Variant effect scores in B73

The variant effect scores are calculated by using the Evolutionary Scale Modeling (ESM1b) protein language model (Z. Lin *et al*., 2023) through the esm-variants tool (Brandes *et al*., 2023). The esm-variants tool calculates variant effect scores based on the log-likelihood ratio difference between the variant and its wild-type version. Scores above -7 indicate benign outcomes, while scores below -7 suggest possible phenotypic effects (Brandes *et al*., 2023). A score of 0 denotes that the variant and wild-type are the same. Over 550 million missense mutations are possible across all isoforms in the B73 proteome. If only the canonical isoforms for each B73 gene model are considered, there are over 273 million possible missense mutations. Approximately 42% of these mutations were predicted as having a possible phenotypic effect (See **Figure 3A** for the distribution of the variant effect scores in the canonical isoforms in the B73 proteome). PanEffect provides a Python script to convert the esm-variants output into a comma-separated (CSV) file containing the x and y position of the heatmap, the variant score, and the amino acid code for the wild-type (B73) and the substitution.

**Figure 3:**
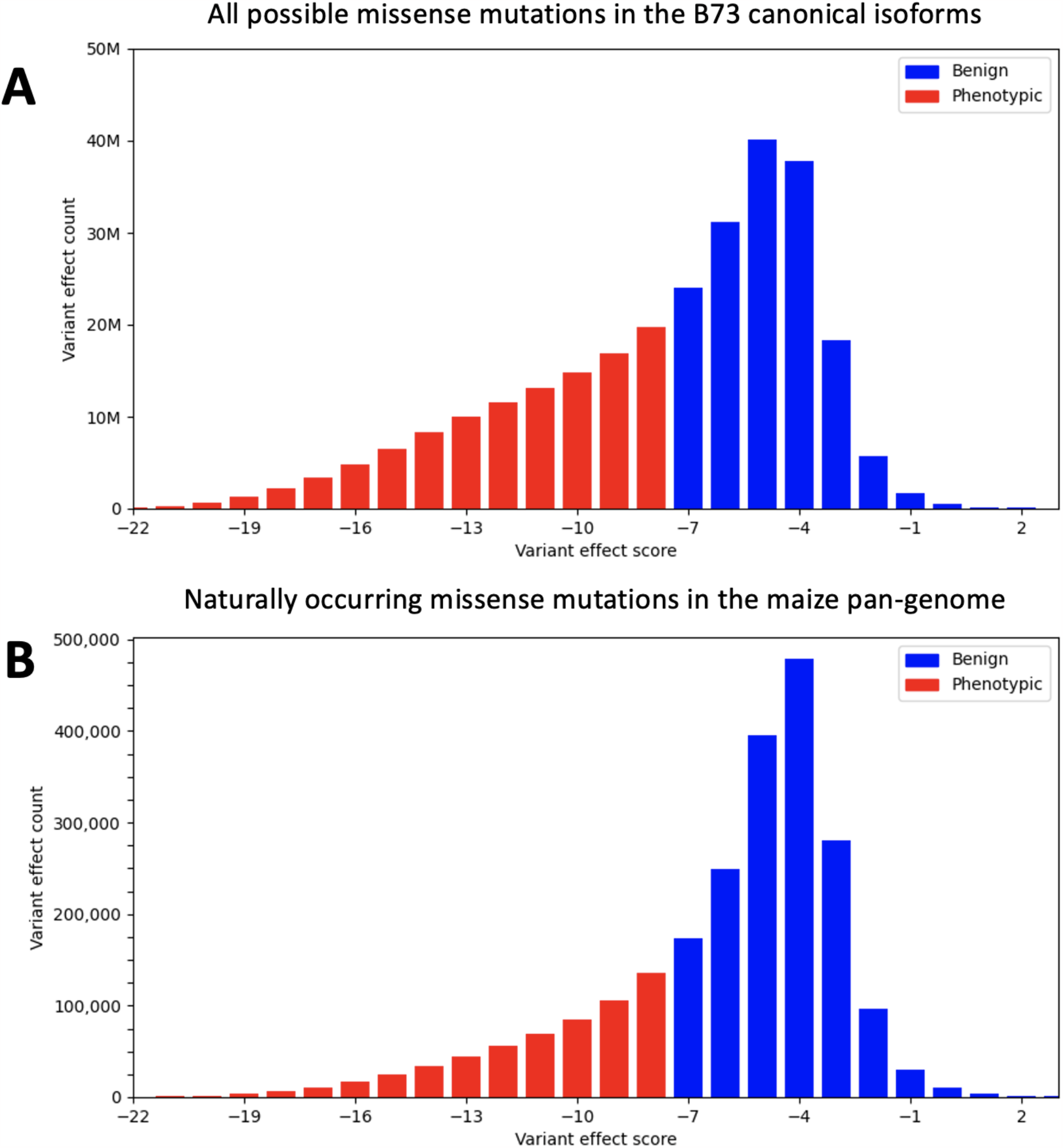
Variant effect scores in B73 and the maize pan-genome. Panel A shows the distribution of the variant effect scores for over 273 million possible missense mutations among the canonical isoforms in the B73 genome. Panel B illustrates the distribution of the variant effect scores of the 2.3 million actual missense mutations between proteins in B73 and 51 other genomes in the maize pan-genome. For both panels, the x-axis is labeled by the variant scores and the y-axis shows the count of variants with that score. Red bars have scores equal to or less than -8 and are considered likely to have a phenotypic effect. The blue bars have scores greater than -8 and are more likely to be benign. Note: the y-axis scale for the B73 plot is tenfold larger, indicating a greater number of possible missense mutations compared to naturally occurring mutations in the pan-genome.

### Pan-genome multiple sequence alignments

The multiple sequence alignments (MSA) were computed using famsa (Deorowicz *et al*., 2016) from pan-genome relationships derived from the software package Pandagma (Cannon, 2023), which uses an all-vs-all approach to align the canonical transcripts in a given pan-gene group. Pandagma generated the MSA on 51 maize reference genomes including two versions of B73 (Hufford *et al*., 2021; Jiao *et al*., 2017), a set of 25 diverse maize lines called the Nested Association Mapping (NAM) panel (Hufford *et al*., 2021), 12 lines used in Chinese breeding programs (Wang *et al*., 2023), 4 European lines (Haberer *et al*., 2020), and an additional set of 8 high-quality maize lines available at MaizeGDB. The maize genome Zm-B73-REFERENCE-NAM-5.0 is used as the reference genome and the B73v5 annotation set contains 39,755 gene models and 75,539 transcripts. The 51 maize genomes are organized and color-coded according to their heterotic group (**Figure 2**).

### Variant effect scores in the pan-genome

The variant effect scores for B73 were combined with the pan-genome multiple sequence alignments to create heatmap representations for all variant effects across the maize pan-genome. Using B73 as a reference, the natural variation in each protein of the pan-genome was scored using B73 as the wild type and the target protein as the variant substitution. Out of the 273 million potential missense mutations in the canonical isoforms, fewer than 1%, approximately 2.3 million, are actually observed within the proteins of the maize pan-genome. Of those coding variations, 74% were predicted as having a benign effect. See **Figure 3B** for the distribution of the naturally occurring variant effect scores in the maize pan-genome. PanEffect provides a Python script to combine these two data types to create a tab-separated (TSV) file listing the x and y positions of the heatmap, the variant score, the positions of the amino acid in B73 and target protein, and the amino acid codes at those positions.

### Protein secondary structures

The three-dimensional protein structures for all protein isoforms in the B73 genome were generated using ESMFold (Z. Lin *et al*., 2023) with the default parameters. These structures are used as input to the DSSP (Define Secondary Structure of Proteins) tool (Kabsch and Sander, 1983) to assign secondary structures (alpha-helices and beta sheets) to each protein. PanEffect has a Python script that converts the DSSP output to a TSV file listing the position, amino acid, and secondary structure code for each B73 isoform.

### Pfam domains

The locations of Pfam functional domains for the B73 proteome were calculated by the NAM sequencing consortium (Hufford *et al*., 2021) using Interproscan (Jones *et al*., 2014) and are available at MaizeGDB. PanEffect provides a Python script to generate a separate TSV file containing the Interproscan information (PfAM position, ID, name, and Gene Ontology terms) for each B73 transcript. An additional Python script is provided to support Pfam searches from Hmmer (Eddy, 2011).

### Functional annotations

MaizeGDB hosts a set of functional annotations for maize genome assemblies including UniProt annotations, canonical gene transcript IDs, and manually annotated gene names and symbols for each gene model in B73 (Cannon *et al*., 2011; Sen *et al*., 2009). The PanEffect search form uses gene models as the default search term, but alternative search terms are supported using a two-column synonym file that links the gene model identifier to an alternative search term. PanEffect has a Python script to parse this information into a TSV file for each B73 gene model.

### GWAS trait annotations

MaizeGDB hosts three GWAS atlas datasets. Each dataset contains sets of single nucleotide polymorphisms (SNPs) linked to specific traits using genome-wide association studies (GWAS). The first set was compiled in 2014 (Wallace *et al*., 2014) and has GWAS mappings for over 40 traits to 40,000 SNP locations in the B73 genome. The second set is from the 2022 update of the GWAS Atlas database (Tian *et al*., 2020), which combined GWAS data from 133 papers covering 531 studies for 279 traits across 42,000 SNP loci. The third dataset is from a 2022 study (Li *et al*., 2022) that performed GWAS for 21 important agronomic traits across 1,604 inbred lines and identified 2,360 significant associations at 1,847 SNP loci. PanEffect has a Python script that links a trait to a gene model by finding any SNP position within 1,000 base pairs of the start and end positions of a B73 gene model. The data for the three datasets are merged, and a final TSV file is created for each gene model listing the trait name and which study it came from.

## Conclusions

The influence of genetic variations on phenotypes is central to plant breeding and genetics. PanEffect can contribute to understanding this relationship by offering a framework to integrate trait and functional annotations with AI-predicted structures and variant effect scores at both reference genome and pan-genome levels. Researchers and breeders can quickly gain insights into the potential consequences of coding variants across numerous maize genomes, allowing them to view the scope of amino acid substitutions in the B73 maize reference genome and discern effects across the maize pan-genome. This is particularly critical as global challenges like climate change and food security become more pressing. PanEffect’s simplicity, utility, and free availability at MaizeGDB and GitHub make it suitable for widespread adoption, customization, and enhancement, providing a valuable alternative to more resource-intensive models like AlphaFold, and promoting accessibility to these approaches.

## Acknowledgments

The work for this project was performed on the Atlas and Ceres high-performance clusters as part of the USDA-ARS SCINet initiative. We would like to thank the SCINet administrative staff and the Virtual Research Support Core team.

## Data availability

PanEffect is a free resource and a maize instance is hosted at https://www.maizegdb.org/effect/maize/. The source code and documentation are available on GitHub at https://github.com/Maize-Genetics-and-Genomics-Database/PanEffect. The full datasets can be downloaded from MaizeGDB at the Artificial Intelligence link: https://www.maizegdb.org/download.

